# Syncytial coupling of mid-capillary pericytes underlies seizure-associated electro-metabolic signaling

**DOI:** 10.64898/2026.03.16.711912

**Authors:** Mirja grote Lambers, Majed Kikhia, Agustin Liotta, Han Wang, Henrike Planert, Thilo Kalbhenn, Ran Xu, Julia Onken, Thomas Sauvigny, Ulrich-W Thomale, Angela M Kaindl, Martin Holtkamp, Pawel Fidzinski, Matthias Simon, Henrik Alle, Jörg RP Geiger, Christian Madry, Richard Kovács

**Affiliations:** Institute of Neurophysiology; Department of Neurology and Experimental Neurology; Einstein Center for Neurosciences; Department of Neurosurgery; Pediatric Neurosurgery; Department of Pediatric Neurology; Center for Chronically Sick Children; Institute of Cell- and Neurobiology; German Epilepsy Center for Children and Adolescents; Department of Clinical and Experimental Epileptology, Charité - Universitätsmedizin Berlin, Germany; Department of Neurosurgery Bethel Clinic - University of Bielefeld Medical Center OWL, Germany; University Medical Center Hamburg-Eppendorf, Germany; DZHK (German Centre for Cardiovascular Research), Berlin, Germany; German Center for Child and Adolescent Health (DZKJ), section CNS development and neurologic disease, partner site Berlin, Germany

**Keywords:** pericytes_1_, epilepsy_2_, potassium signaling_3_, neurovascular coupling_4_, slice cultures_5_, neurovascular unit_6_

## Abstract

Disturbances of neurovascular coupling (NVC) contribute to metabolic derailment and neurological symptoms associated with epilepsy. While postictal arterial constriction can be alleviated by inhibitors of voltage gated calcium channels (VGCCs), less is known regarding seizure-associated electrical signals in higher-order capillaries and their role in determining pericyte tone during seizures.

Here we investigated electrical signaling within the ex vivo neurovascular unit (NVU) derived from rat and human brain tissue. We focused on electrical signal transduction between pericytes and endothelial cells and the potential role of VGCCs in vasomotion. Using dye coupling and paired patch-clamp recordings, we showed that morphologically heterogeneous groups of mid-capillary pericytes build a functional syncytium with endothelial cells. Coupling was asymmetric, allowing for directed propagation of electrical signals.

Regardless of their morphology, mid-capillary pericytes responded with depolarization and constriction to metabotropic receptor (GPCR) activation (by thromboxane, norepinephrine and UDP-glucose). However, depolarization via the patch pipette induced neither Ca^2+^-influx nor constriction, suggesting lack of contribution of VGCCs to vasomotion.

On inducing epileptiform activity, A2a adenosine receptors and inwardly rectifying potassium channels hyperpolarized the capillary syncytium, followed by repeated depolarizations due to seizure-associated potassium increase in the parenchyma. Thus, while mid-capillary pericytes are contractile, their tone does not rely on their membrane potential and VGCCs. However, syncytial coupling allows for transmission of seizure-associated hyper- and depolarizing signals to upstream feeding arterioles.

## INTRODUCTION

Efforts to identify novel targets for preventing epileptogenesis have increasingly focused on the neurovascular unit (NVU). Disturbances in neurovascular coupling and capillary hypoperfusion contribute to epilepsy-associated cognitive disturbances [1] [2] [3] [4]. Capillary pericytes play a pivotal role in orchestrating the hemodynamic response to increased energy demand during neuronal activity. Pericyte injury contributes to seizure-induced neurovascular uncoupling [5] [6], pathological vascular reorganization [7] [8] as well as to dysfunction of the blood-brain barrier (BBB). The latter has been implicated in inducing perivascular inflammation and pro-epileptogenic transformation of the NVU [9] [10] [11] [12] [13].

Coordination of the hemodynamic response necessitates long range communication along the vasculature. The membrane potential of the electrically coupled components of the NVU may transmit metabolism-related signals to upstream feeding arterioles, a process referred to as electro-metabolic signaling [14] [15] [16]. Key determinant of the metabolism-dependent electrical signals is the neuronal activity-associated increase in parenchymal potassium concentration [17] [18]. Potassium released from the astrocytic endfeet facing endothelial and smooth muscle cells (SMCs) contribute to cerebral functional hyperemia [19] [20] [21] [22] [23] by enhancing open channel conductance of endothelial Kir2.1 potassium channels [24] [25] [26]. KATP channels, consisting of the pericytic Kir6.1 channel and the accessory sulfonylurea receptor SUR2, function as a metabolic sentinel, linking an increase in the ADP/ATP ratio to hyperpolarization of the pericyte membrane and capillary vasodilation [27] [28] [29]. However, whether this metabolism-dependent hyperpolarization directly influences capillary pericyte tone or acts indirectly by modulating Ca^2+^ signaling in upstream arteriolar SMCs, remains unclear.

Cerebral pericytes represent a diverse group of mural cells, distinguished by their embryonic origin, protein expression, morphology (e.g., ensheathing, mesh, and thin-strand pericytes), and position along the capillary bed, each likely serving unique roles in blood flow regulation and BBB maintenance [30] [31] [32] [33] [34] [35] [36] [37] [38] [39] [40] [41]. Ensheathing pericytes of the post-arteriolar transition zone are contractile, expressing alpha-smooth muscle actin (α-SMA) and voltage-gated Ca^2+^ channels (VGCCs). It is now widely recognized that mid-capillary mesh and thin-strand pericytes are capable to regulate capillary diameter [32] [42] [6] [27], but little is known about the mechanisms behind the slow capillary vasomotion. Optogenetically induced depolarization in channelrhodopsin-expressing thin-strand pericytes triggered vasoconstriction [43]. Spontaneous or intraluminal pressure induced Ca^2+^ oscillations in thin-strand pericytes were dependent on extracellular Ca^2+^ influx in part via VGCCs [44] [45]. However, direct in situ electrophysiological evidence on the membrane potential dependence of capillary tone is scarce [46] and even less is known about the capillary electro-metabolic signaling during recurrent seizures. We have previously described seizure -dependent inward membrane currents in rat capillary pericytes [6] but it remained uncertain what this means for the capillary tone and how these findings can be translated into the human context.

Here we investigated seizure-associated membrane potential changes in capillary endothelial cells, various morphological classes of pericytes and astrocytes using a combination of fluorescence microscopy, potassium ion-sensitive electrodes, and whole-cell patch-clamp recordings. We characterized the capillary syncytium by examining diffusion of a fluorescent tracer and electrical signaling between pericyte-pericyte and pericyte-endothelial cell pairs in organotypic hippocampal slice cultures (OHSCs) and acute hippocampus slices. Furthermore, capillary pericyte reaction to pharmacologically induced epileptiform activity was confirmed in human brain slices obtained from epilepsy surgeries.

## RESULTS

### 1. Mid-capillary pericytes represent a morphologically and electrophysiologically heterogeneous group

We classified capillary pericytes in OHSCs based on their morphology and electrophysiological properties. Pericytes were identified prior to recordings by labeling with MitoSox [47] [6] or NeuroTrace ([48]; Figure 1A, suppl. movie 1). As both probes preferentially label nuclei and organelles, the morphological classification was performed using a second fluorescent dye (OGB or rhod-2, respectively) introduced via the patch pipette. Rat capillary pericytes (n = 336 selected out of 352 total, based on imaging quality criteria), could be divided into four classes based on morphological properties by using unsupervised clustering (Figure 1B). The classes were i.) elongated thin-strand pericytes with very few processes, the main shank often encircling the capillary, ii.) mesh pericytes with numerous circumferential processes enwrapping the capillary, iii.) transitional pericytes with intermediate features between the first two types, and iv.) short pericytes with mossy processes eventually resembling postcapillary pericytes (Figure 1. B-E).

**FIGURE 1.**
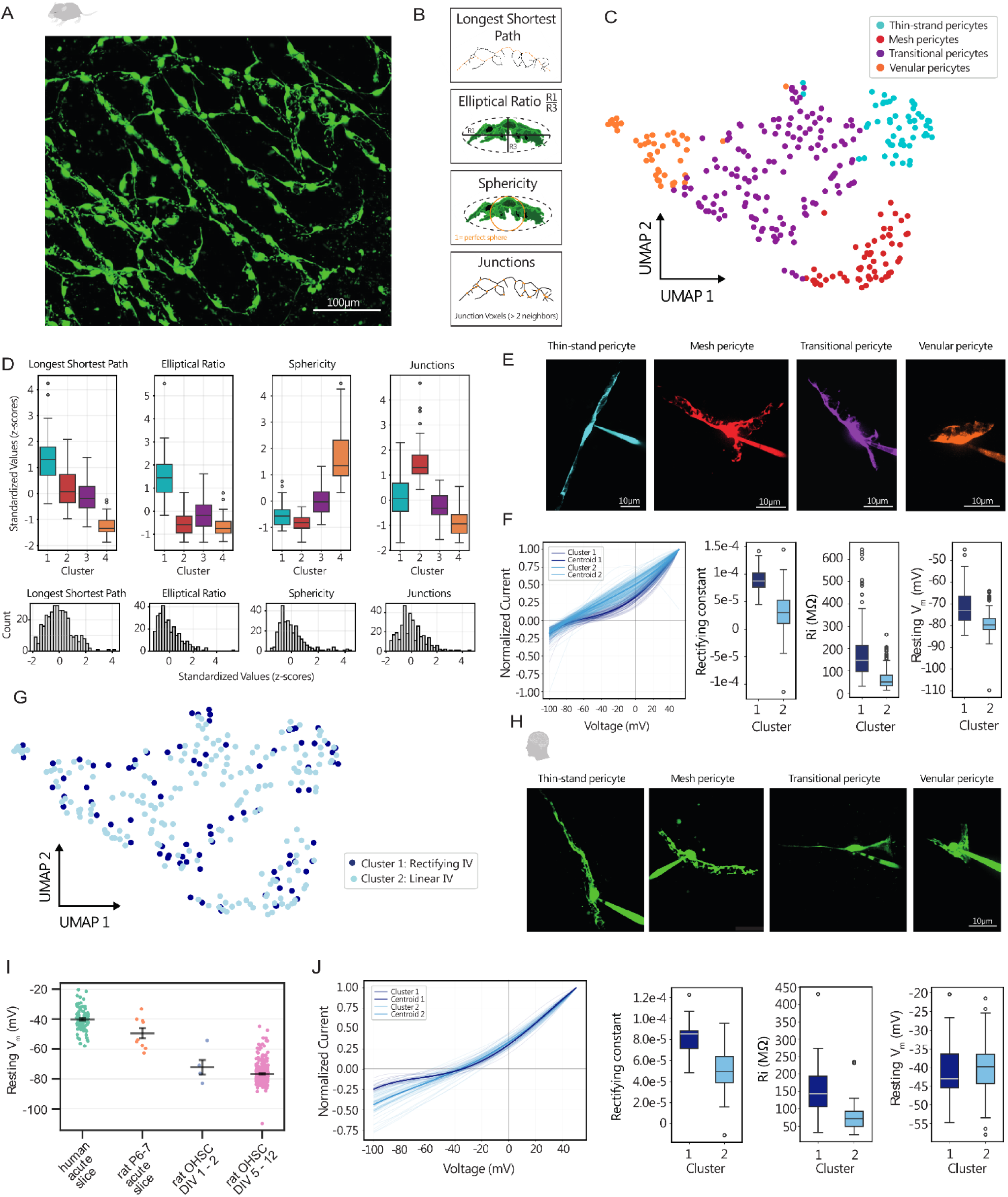
Pericyte phenotypes in organotypic slice cultures (OHSC) and human brain slices. **(A)** Pericyte network in an OHSC stained with NeuroTrace 500/525. **(B)** Based on silhouette scores we selected four morphological parameters for clustering individual pericytes following patch clamp recordings: longest shortest path, sphericity, elliptical ratio and number of junctions (see methods). **(C)** UMAP of pericyte clusters based on morphology (4 clusters by using KMeansWithNulls, n = 336). Each point corresponds to a pericyte. The colors of the clusters correspond to the colors in D and E. **(D)** Distribution of parameters within the four clusters. Boxes showing the 25th to 75th percentile, the bar represents the median value of the distribution and whiskers extend to the most extreme values within 1.5 × interquartile range from the quartiles. Cluster 1 contains elongated thin-strand pericytes, cluster 2 is characterized by mesh-pericytes with numerous junctions, cluster 3 comprises shorter transitional pericytes and cluster 4 contains short, spherical cells. Be low: Histograms of parameter distribution of all 336 cells included. **(E)** Representative fluorescence images of pericytes from each cluster. **(F)** Clustering based on electrophysiological properties revealed two groups (dark blue and light blue, darker lines show the centroids of the clusters) with different membrane potential (Vm), input resistance (Ri) and current-voltage relationships (Cluster 1 (dark blue): Ri = 198.99 ± 15.37 MΩ, Vm = -70.97 ± 0.94 mV, n = 94; Cluster 2 (light blue): Ri = 64.15 ± 2.8 MΩ, Vm = -78.87 ± 0.34 mV, n = 229), cluster 1 showing stronger rectification than cluster 2. **(G)** Mapping the electrophysiological clusters onto the morphological UMAP reveals no clear distinction between the two features. **(H)** Representative fluorescence images of human pericytes show similar morphological characteristics as in OHSCs. **(I)** The resting membrane potential of human pericytes in acute slices (Vm = -40.18 ± 0.76 mV, p^ = 0.81 [0.76, 0.85], n = 93) was slightly more positive than that of acute rat pericytes (Vm = -49.5 ± 3.48 mV, p^ = 0.66 [0.59, 0.72], n = 9, p= 0.145). During the culturing process the membrane potential of the pericytes in acute slices becomes progressively more negative (DIV 1-2: Vm = -72.23 ± 4.85 mV, p^ = 0.31 [0.23, 0.41], n = 5, p = 0.002; DIV 5-12: Vm = -76.62 ± 0.41 mV, p^ = 0.22 [0.16, 0.28], n = 309, p < 0.001). **(J)** Clustering of human pericytes (n = 93) based on electrophysiological parameters revealed 2 groups (Cluster 1 (dark blue): Ri = 171.53 ± 24.28 MΩ, Vm = -40.64 ± 2.00 mV, n = 19; Cluster 2 (light blue): Ri = 75.48 ± 4.30 MΩ, Vm = -40.10 ± 0.81 mV, n = 74). Pericytes in cluster 1 showed higher input resistance (Ri) and increased rectification, similar to group 1 pericytes in OHSCs.

Whole-cell patch clamp recordings using low-chloride (8 mM) intracellular solution revealed highly negative membrane potentials (Vm) in pericytes -78.86 ± 0.70 mV, n = 75) comparable to the values obtained in astrocytic endfeet (−79.85 ± 0.88 mV, n = 48). Using intracellular solutions with more physiological intracellular chloride concentration (34 mM, [49]), Vm values remained similar -75.90 ± 0.49 mV, n = 234), indicating limited influence of chloride conductance in resting pericytes in OHSCs. While astrocytes exhibited low input resistance (Ri) and a stereotypic linear current-voltage (I-V) relationship, pericytes showed greater heterogeneity in their electrophysiological properties. Unsupervised K-Means clustering divided the recorded pericytes into two groups, i.) the majority of the pericytes with low input resistance and an ohmic I-V profiles (Ri: 64.15 ± 2.80 MΩ, Vm: -78.87 ± 0.34 mV), and ii) a smaller fraction presented with higher input resistance, more positive Vm, and rectifying I-V relationship (Ri: 198.99 ± 15.37 MΩ, Vm: -70.97 ± 0.94 mV, Figure 1F). Notably, no correlation was found between the electrophysiological and morphological classification (Figure 1G). Resting membrane potential obtained in endothelial cell recordings (Vm: -80.95 ± 1.03 mV, n = 14) and input resistance (64.58 ± 7.75 MΩ, n = 14) was in the range of the first type of pericytes.

We then investigated the morphological and electrophysiological properties of pericytes in acute human cortex slices. Pericytes were selected at capillaries with a diameter <10 µm and the identity of the cells was confirmed by post hoc morphological validation after dye loading and/or by the presence of coupling to other cells of capillary syncytium (see below). Despite size differences, human pericytes fell into the same categories as rat pericytes (thin-strand, mesh and transitional) (Figure 1H). Remarkably, the resting Vm of human pericytes in acute slices (−40.18 ± 0.76 mV, n = 93) was more positive than rat pericytes in slice cultures. To determine whether this depolarization was species-specific or related to the type of preparation, we conducted whole-cell recordings in acute slices from juvenile rats (P6–7) shortly after slicing, and during the first two days in culture (DIV 1–2). Median pericytic Vm was -49.50 ± 3.48 mV (n = 9) immediately (>2 hrs) after slicing in juvenile rats, slightly more hyperpolarized than that recorded in adult human tissue. Upon culturing, Vm gradually decreased to -72.23 ± 4.85 mV (n = 5) within 48 hours, and to -76.62 ± 0.41 mV (n = 309) after 5 days in vitro (Figure 1I). Notably, the positive Vm in acute slices was accompanied by a vasoconstriction, while capillaries in culture were dilated, as described previously ([6]). The similarities between juvenile rats and adult human acute slices indicate that Vm is most likely dependenton the preparation (acute vs culture) and not on the species. Clustering of human pericytes based on electrophysiological parameters identified both linear and rectifying types as seen in rat slice cultures (Figure 1J). In conclusion, the morphological diversity of mid-capillary pericytes - both rat and human - is not reflected in their electrophysiological properties.

### 2. Pericytes and endothelial cells constitute an electrical syncytium

Electrical coupling to other components of the NVU might influence the electrophysiological properties of pericytes. To address this, we next characterized gap junctional coupling characteristics underlying the functional syncytium of rat and human capillaries. By using a pipette solution with the gap-junction permeable AlexaFluor488 (n = 27) we observed diffusion of the fluorescent probe from the recorded pericytes into endothelial cells and other pericytes up to ∼500 µm distance crossing several capillary bifurcations (Figure 2A). Labeling the endothelial cell adjacent to the recorded pericyte always preceded staining of neighboring pericytes along the capillary, whereas astrocytes - despite the proximity of the endfeet to the capillary wall - were never labelled. On the contrary, recordings of astrocytes with AlexaFluor488 containing solution (Figure 2B, n = 8) revealed dye diffusion within the astrocytic syncytium but not to endothelial cells or pericytes. These findings indicate that endothelial cells and pericytes on the one side and astrocytes on the other side form two separate functional syncytia (Figure 2AB). Input resistance of pericytes was inversely correlated (Spearman-Rho -0.68, p = 0.00005, Figure 2CDa) to the coupling index, defined as the ratio of fluorescence intensity at the contact site between the recorded cell and the soma of the subsequently labeled cell. Gap junctional coupling was also present between pericytes and endothelial cells in the human capillaries (Figure 2 Db). Similarly to OHSCs, coupling index was the highest between the recorded pericyte and the adjacent endothelial cell, whereas in endothelial cell recordings dye coupling to another endothelial cell was stronger than to pericytes of the same capillary. To further investigate the electrophysiological implications of the syncytial organization we obtained paired patch clamp recordings from pericyte-pericyte and pericyte-endothelial cell couples both in OHSCs and in human cortex slices. Recorded cells were identified by using different AlexaFluor probes in the pipette solution. To describe the electrical coupling, we used the ratio of membrane voltage changes between the recipient and stimulated cells in response to brief hyper- and depolarizing current steps (coupling coefficient, Figure 2F). A coefficient near 1 indicates near-lossless electrical coupling, whereas 0 reflects complete electrical isolation. The membrane potential did not differ significantly between endothelial cell-pericyte couples (ΔVm: -0.49 ± 4.14 mV, n = 7, p^ = 0.55 [0.1, 1.0], p = 0.8). In pericyte-pericyte pairs, the difference was borderline significant (ΔVm: 5.06 ± 2.04 mV, n = 10, p^ = 0.34 [0.183, 0.5], p = 0.05). In cell pairs with a high coupling coefficient occasional spontaneous membrane potential oscillations were synchronized (Figure 2G). Remarkably, not only were resting Vm values nearly identical, but both cells also exhibited similar I–V profiles, regardless of their morphological classification. This was also observed in endothelial cell and pericyte paired recordings, suggesting that the electrophysiological properties recorded from individual cells may reflect the behavior of the capillary syncytium as a whole.

**FIGURE 2.**
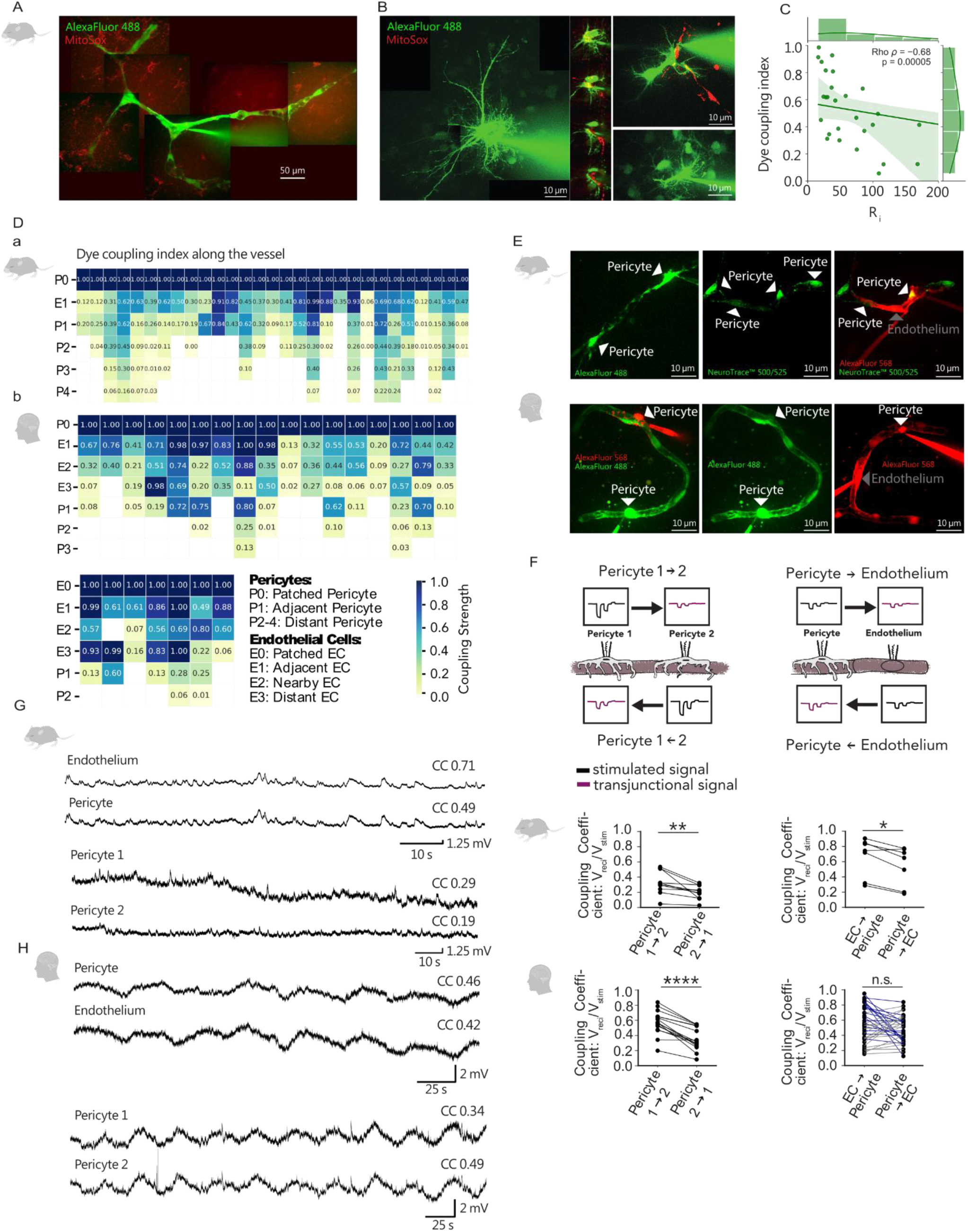
Pericytes and endothelial cells form an electrically coupled syncytium. **(A)** Spread of AlexaFluor 488 fluorescent dye along the capillaries reveals bidirectional coupling of pericytes and endothelial cells via gap junctions in MitoSox labeled OHSCs. **(B)** Images showing the astrocytic syncytium in OHSC visualized by patching an astrocyte with Alexa488 in the pipette (n = 8). The astrocytic syncytium (green channel, Alexa Fluor 488) i s not connected to the pericytes (red channel, MitoSox), which are located directly below the astrocyte endfeet. The images show three different examples. Note the subsequently labelled astrocytes in the surrounding. The excerpts on the left side of the first image represent different z -planes from the point of contact between pericytes and astrocytes. **(C)** The coupling index is calculated as the ratio of fluorescence intensity of the recorded cell processes in relation to the soma of the subsequently stained cells. Input resistance showed an inverse correlation with the coupling index (ρ = -0.68, p = p < 0.0001) in pericytes. **(D)** Dye coupling matrix of 27 rat **(a)** and 17 human **(b)** pericytes and 7 endothelial cells, color-coded from blue to yellow, indicating varying degrees of coupling strength . The absence of dye in the surrounding astrocytic endfeet ruled out non-specific dye leakage. **(E)** Examples of dye coupling in paired recordings in OHSC (top) and human temporal lobe tissue (bottom) patched with Alexa568 or Alexa488 dye. Upper images from recordings in OHSCs. On the left: Alexa488 stained p ericyte-pericyte couple, middle: NeuroTrace labeled capillary segment, right: pericyte-endothelial cell couple recorded with Alexa568 in the same segment. Bottom images from recordings in human slices. On the left: pericyte-pericyte couple with different Alexa probes (488/568), middle: same capillary only with the green channel presented to illustrate diffusion of Alexa488 between the two cells with no apparent contact, right: pericyte -endothelial cell couple with Alexa568. Scale bars: 10 µm. **(F)** Simultaneous dual patch-clamp recordings of pericyte-endothelium or pericyte-pericyte pairs. Upper diagrams illustrate the experimental setup of measuring electrical coupling: stimulated signals are given in black and transjunctional responses in purple. Quantification of the coupling coefficients for pericyte -to-pericyte connections and pericyte-to-endothelium connections showing asymmetrical coupling in OHSC (top) and human tissue (bottom) . OHSC pericyte-pericyte coupling: n = 10, p^ = 0.22 [0.07, 0.37], p = 0.002; OHSC pericyte-pericyte coupling: n = 7, p^ = 0.29 [0.13, 0.44], p = 0.03; human acute slice pericyte-pericyte coupling: n = 16, p^ = 0.15, [0.04, 0.26], p < 0.0001; human acute slice endothelial-pericyte coupling: p^ = 0.44 [0.31, 0.57], p = 0.36). **(G)** Representative traces of an endothelial-pericyte and pericyte-pericyte pair in OHSC, displaying synchronous potential changes for strongly coupled cells in contrast to weakly coupled cells. **(H)** Representative traces of an endothelial-pericyte and pericyte-pericyte pair in human tissue.

Electrical coupling between connected cell pairs was asymmetricalin both rats and humans. In OHSCs, the coupling coefficient from endothelial cells to pericytes was higher than between pericyte–pericyte pairs, and the coupling often exhibited directionality towards the pericyte. Pericyte to pericyte coupling also showed significant directionality in most of the recorded pairs. Interestingly, electrical coupling was observed between pericytes that apparently lacked any direct contact between their processes, suggesting that communication may occur indirectly via the endothelial syncytium (Figure 2E). The lowest coupling coefficients were recorded in pericyte pairs separated by a capillary bifurcation, implying that these junctions may act as barriers to the propagation of electrical signals. Consistent with findings showing dye diffusion along the human capillary syncytium, pericyte–pericyte and pericyte–endothelial cell pairs in human cortex slices also exhibited electrical coupling. The coupling coefficient between human pericytes was higher than in rat tissue (Figure 2F), and correspondingly, the difference between the resting Vm was smaller (endothelial cell-pericyte ΔVm: - 0.59 ± 0.44 mV, n = 35, p^ = 0.52 [0.49, 0.55], p = 0.19; pericyte-pericyte ΔVm: -2.10 ± 1.60 mV, n = 16, p^ = 0.56 [0.47, 0.66], p = 0.27). Although human endothelial-to-pericyte coupling was also asymmetric, the direction varied: approximately half of the cell pairs exhibited signal bias toward the endothelial cell. In conclusion, our data demonstrates the presence of a capillary electrical syncytium comprising endothelial cells and pericytes while the observed asymmetry in electrical coupling is well-suited to support directional signal transmission.

### 3. Syncytial electrical signals and vasomotion in OHSCs

Electro-metabolic signaling for capillaries would necessitate contractility of mid-capillary pericytes and the presence of voltage dependent Ca^2+^ influx, to couple membrane depolarization with increasing vasotonus [14]. Regarding contractility, we previously found that mid-capillary thin-strand and mesh pericytes constricted reversibly in response to mechanical forces associated with establishing the whole-cell configuration [6]. Additionally, application of the stable thromboxane analogue U46619 constricted capillaries [35], which could be reversed by NO or by recurrent seizure activity [6].

Since contractility is present in mid-capillary pericytes in OHSCs, we further investigated whether membrane depolarization is necessary for the vasoconstriction induced by various GPCR agonists. Norepinephrine [50] or the P2Y14 receptor agonist UDP-glucose both resulted in depolarization and vasoconstriction (Figure 3A). Despite the continued presence of the agonists, depolarization and capillary constriction were only transient. In contrast, application of U46619 resulted in a sustained depolarization and vasoconstriction (Figure 3B and F, suppl. movie 2). In human pericytes, U46619 resulted in a further depolarization by an additional 16.71 ± 3.25 mV (n = 3) from the already positive resting membrane potential.

**FIGURE 3.**
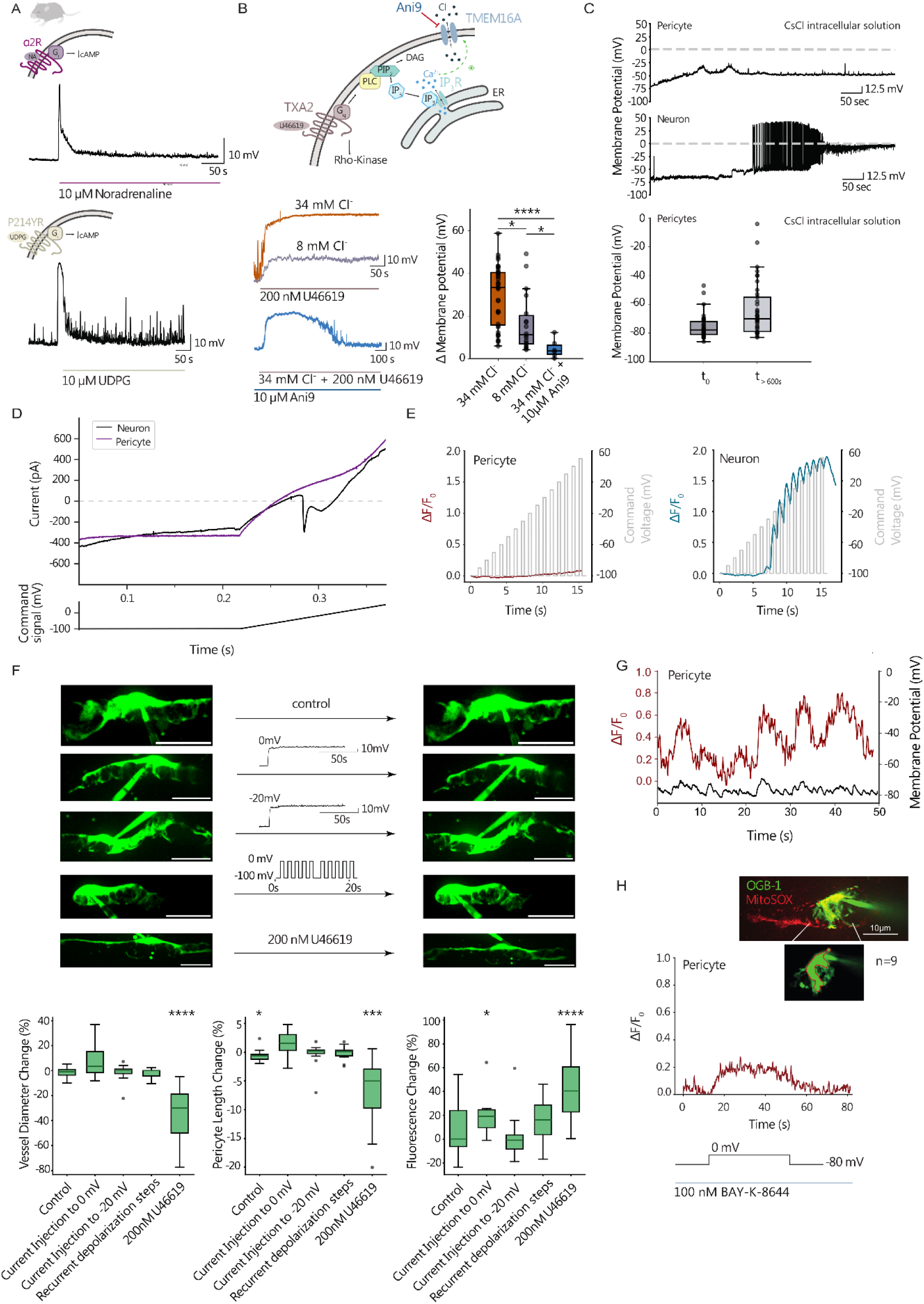
Depolarization and constriction are not interdependent in mid-capillary pericytes. **(A)** Changes in pericyte membrane potential after norepinephrine and UDP-glucose application. **(B)** The thromboxane analogue U46619 depolarizes and constricts pericytes. Upper left corner: The amplitude of the depolarization induced by U46619 depended on the intracellular chloride concentration. See the orange trace (34 mM Cl⁻) and the light purple trace (8 mM Cl⁻). Lower left and right panel: The plateau phase was reversed by the TMEM16 inhibitor Ani9 (blue trace), indicating the involvement of Ca^2+^-activated Cl^-^ channels. Significant difference between 34 mM Cl⁻ and 34 mM Cl⁻ + 10 µM Ani9 (p = 0.0001), between 8 mM Cl⁻ and 34 mM Cl⁻ (p = 0.03) as well as between 8 mM Cl⁻ and 34 mM Cl⁻ + 10 µM Ani9 (p = 0.02); 34 mM Cl⁻: n = 39, p^ = 0.74 [0.69, 0.79]; 8 mM Cl⁻ : n = 18, p^ = 0.53 [0.46, 0.61]; 10 µM Ani9: n = 6, p^ = 0.22 [0.16, 0.30]. **(C)** Upper panel: Recording of the resting membrane potential (Vm) of a pericyte and a neuron immediately after establishing the whole cell configuration with a CsCl-containing intracellular solution. In contrast to neurons, pericytic depolarization is moderate. Lower panel: Quantification of pericyte membrane potential immediately after rupturing the membrane and equilibration with the CsCl intracellular solution. Although the difference reached statistical significance (p^ = 0.73 [0.65, 0.81], p < 0.001), the absolute change was modest (t0: −75.68 ± 1.37 mV; t1: −63.51 ± 3.12 mV, mean ± SEM, n = 37), remaining far from the strong depolarization typically observed in neurons patched with CsCl-based intracellular solution. **(D)** Representative recording of a voltage ramp (−100 mV to 60 mV) of a neuron (turquoise) and pericyte (red), indicating the presence of voltage gated inward currents in neurons but not in pericytes. **(E)** Relative fluorescence change of OGB-1 in response to depolarization steps starting from a holding potential of -100 mV. Neurons showed a marked increase in fluorescence for steps above -50 mV while pericytes solely presented with continuous slight baseline increase. **(F)** Comparison of the percentage change in vessel diameter, pericyte length, and fluorescence change upon i. control (n = 11, vessel diameter: p^ = 0.44 [0.35, 0.53], p = 0.23; pericyte length: p^ = 0.45 [0.4, 0.5], p = 0.05; rhod-2 fluorescence: p^ = 0.52 [0.47, 0.57], p = 0.41), ii. different depolarizing current injection protocols (depolarization to 0 mV (n = 6, vessel diameter: p^ = 0.58 [0.32, 0.85], p = 0.28; pericyte length: p^ = 0.53 [0.46, 0.6], p= 0.22; rhod-2 fluorescence: p^ = 0.64 [0.5, 0.78], p = 0.03), depolarization to -20 mV (n = 13, vessel diameter: p^ = 0.52 [0.46, 0.58], p = 0.51; pericyte length: p^ = 0.50 [0.47, 0.53], p = 0.69; rhod-2 fluorescence: p^ = 0.50 [0.42, 0.58], p = 0.96), recurrent depolarization steps (n = 11, vessel diameter: p^ = 0.48 [0.45, 0.50], p = 0.19; pericyte length: p^ = 0.51 [0.46, 0.56], p = 0.55; rhod-2 fluorescence: p^= 0.56 [0.47, 0.66], p = 0.19) or iii. 200 nM U46619 (n = 14, vessel diameter: p^ = 0.27 [0.16, 0.37], p < 0.0001; pericyte length: p^ = 0.42 [0.38, 0.47], p = 0.001; rhod-2 fluorescence: p^ = 0.63 [0.58, 0.68], p < 0.0001). Vessel diameter, pericyte length and rhod-2 fluorescence changed significantly upon U46619 (vessel diameter change: -33.92 ± 5.54 %; pericyte length change: -7.15 ± 1.61 %; rhod-2 fluorescence increase: 42.77 ± 7.33 %) in contrast to depolarization protocols. Scale bar: 10 µm. **(G)** Representative recording of a pericyte exhibiting spontaneous Ca2+ fluctuations in the presence of the VGCC activator BAY-K-8644 (100 nM). **(H)** Change in OGB-1 fluorescence upon depolarization of the pericyte (on the excerpt) to 0 mV under BAY-K-8644. Slow increase in fluorescence was observed in 7 out of 9 cells.

At physiological intracellular Cl^-^ concentration the amplitude of the depolarization was significantly higher than with low Cl^-^, reaching the calculated reversal potential in ∼40 % (n = 71) of the recorded cells. Furthermore, inhibition of Ca^2+^-activated Cl^-^ channels (CaCCs) by Ani9 reversed the second phase of U46619 induced depolarization, further indicating the involvement of these channels upon GPCR mediated Ca^2+^-release from the intracellular stores (Figure 3B; n = 6).

As CaCCs sufficiently depolarize the pericytes to potentially open VGCCs [27], we next investigated whether the latter are present on mid-capillary pericytes. Whole-cell recordings were performed on pericytes and neurons using 135 mM CsCl containing pipette solutions to block potassium channels and unmask the contribution of VGCCs. Neurons depolarized to approximately 0 mV within minutes after establishing the whole cell configuration in the presence of Cs^+^ and adepolarizing ramp protocol (−100 to +50 mV) revealed the presence of inward Na^+^ and Ca^2+^ currents (Figure 3CD). In contrast, pericytes depolarized more slowly, reaching a steady membrane potential of −63.51 ± 3.12 mV after 10 min, indicating that Cs^+^ cannot block all Kir channels in the syncytium. Neither depolarizing voltage steps nor the ramp protocol from -100 to +50 mV could reveal voltage gated inward currents (Figure 3CD), suggesting lack of VGCCs in pericytes in slice cultures.

In addition to the current recordings in CsCl intracellular solution we conducted Ca^2+^ imaging with different fluorescence probes (rhod-2, OGB-1) depending on the labeling used to identify pericytes (NeuroTrace / MitoSox). Passage of these probes via gap junctions was virtually absent and the fluorescence remained confined to the recorded cell. Depolarizing steps (-100 to +50 mV), which reliably triggered dendritic Ca²⁺ signals in neurons failed to elicit significant fluorescence increase in pericytes—regardless of the fluorescence probe used (Figure 3E). In contrast, U46619 application significantly increased OGB-1 fluorescence by 42.77 ± 7.33 % (p < 0.001). Finally, we assessed changes in pericyte length and capillary diameter upon different depolarizing protocols and compared them with the effect of U46619 (Figure 3F). U46619 induced pericyte shortening (-7.15 ± 1.6 %, p = 0.001) and capillary constriction (diameter change at the pericytic site: -33.92 ± 5.5 %, p < 0.001), indicating that Ca²⁺ release from internal stores is sufficient to drive vasoconstriction. In contrast, none of the depolarizing protocols did induce significant changes in intracellular Ca^2+^, capillary constriction or shortening of the pericyte (Figure 3F). However, depolarizing pericytes in the presence of an L-type VGCC activator BAY-K-8644 (100 nM, [45]) led to a gradual increase in [Ca^2+^]i in 7 out of 9 cells (Figure 3GH). Thus, while BAY-K-8644 suggested the presence of VGGCs, the absence of significant Ca^2+^ influx and vasoconstriction upon depolarization indicates that VGCCs do not contribute to the observed vasomotor responses in mid-capillary pericytes.

### 4. Extracellular potassium dependent membrane potential changes in the pericytic endothelial cell syncytium during seizures

As GPCRs were key regulators of the membrane potential of the capillary syncytium, next we sought to determine the membrane potential changes associated with epileptiform activity. Recurrent seizure-like activity induced by the non-selective voltage gated potassium channel blocker 4AP (100 µM) resulted in an elevation of the extracellular potassium concentration from the baseline of 3 mM up to approximately 8 mM during the initial phase of the seizure followed by an undershoot in OHSCs (Figure 4AB, [51]). This seizure-associated extracellular potassium accumulation translated into membrane potential changes in astrocytes and in the pericytic endothelial syncytium (Figure 4CE). The peak depolarization in astrocytes and pericytes was not different from the potential recorded with an ion-sensitive electrode and was unaffected by differences in intracellular Cl^-^ concentration in pericytes. The median depolarizations in pericytes and astrocytes were not different from the depolarization recorded by the potassium ion-sensitive electrodes, indicating that the shift of the membrane potential primarily reflects changes in the Nernst potential for potassium (Figure 4A-C). In simultaneous intracellular and field potential recordings each burst of the seizure -like activity was accompanied by a pericyte depolarization, similar to the ion-sensitive electrode recordings (Figure 4D). Despite the lack of electrical coupling, both the amplitude and kinetics of seizure -associated changes in astrocytic membrane potential were similar to the pericyte-endothelial cell syncytium. Inducing seizure-like activity composed of tonic and clonic phases is notoriously challenging in human brain slices under submerged conditions. We succeeded by using a double perfusion chamber with enhanced oxygenation [52] and by applying bicuculline in low Mg^2+^ and 5 mM K^+^ aCSF. Under these conditions human pericytes (n = 3) displayed potassium dependent depolarizations followed by an undershoot, similar to the recordings in OHSCs (Figure 4E). The individual depolarizations in the recorded pericytes were smaller in human cortex than those in hippocampal slice cultures, likely reflecting lower potassium increases described previously for the human brain slices [53]. Thus, under epileptic conditions the seizure-associated increase in extracellular potassium concentration is the major determinant of the membrane potential of the pericyte - endothelial cell syncytium both in human and rat.

**FIGURE 4.**
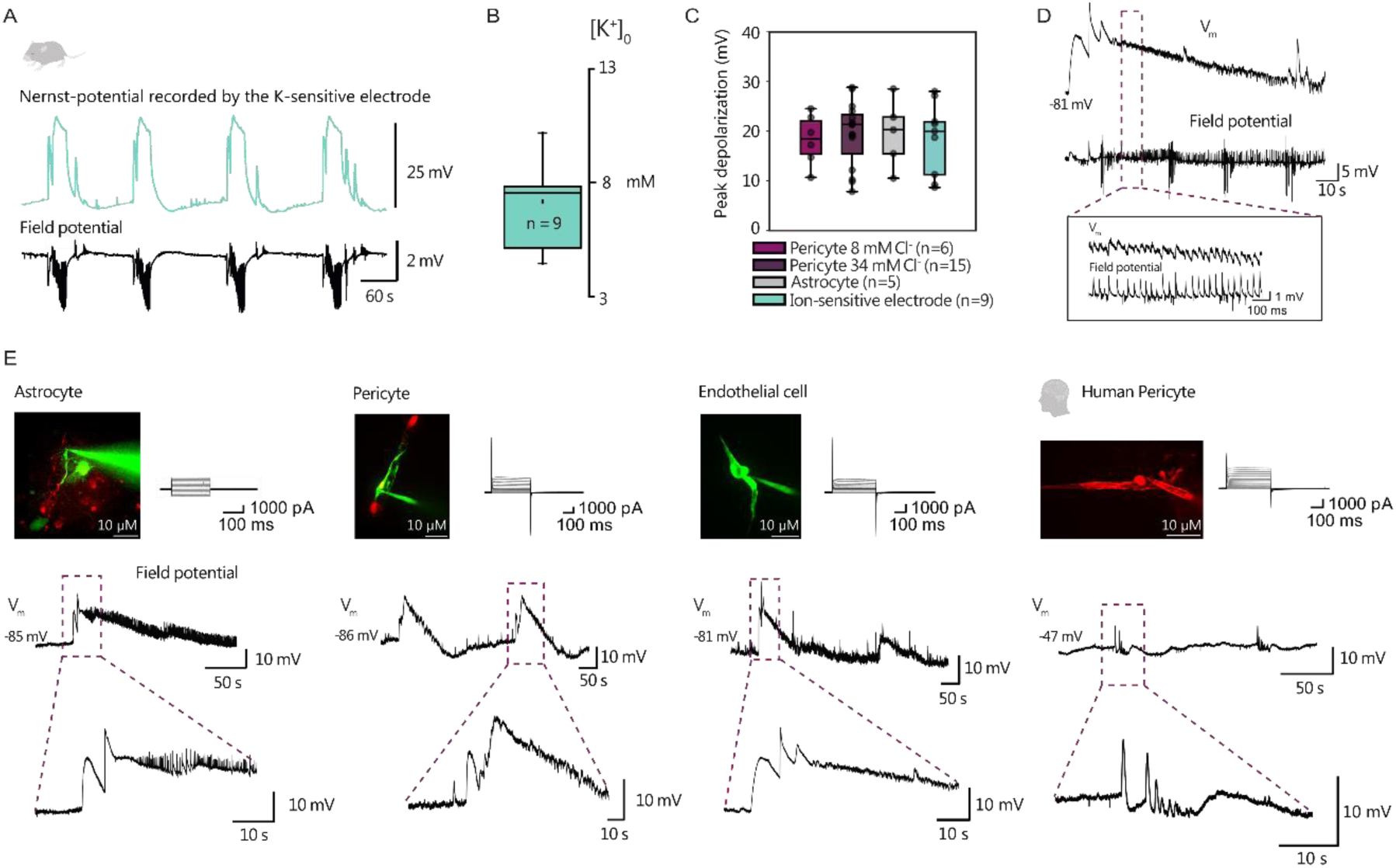
Seizures result in an increase in [K^+^]0, which depolarizes pericytes, endothelial cells and astrocytes. **(A)** Representative recording of a potassium ion-sensitive electrode and concurrent local field potential during recurrent seizures induced by 4AP (100 μM). **(B)** [K^+^]0 increased from 3 to 7.16 ± 0.68 mM (n = 9) during the tonic phase of a seizure. **(C)** Comparison of the changes in membrane potential and ion-sensitive recordings revealed that both astrocyte and pericyte Vm passively follow changes in potassium Nernst potential. Pericyte (Low Cl^-^): Mean ± SEM: 18.3 ± 2.1 mV, Median: 18.5 mV, n = 6; Pericyte (High Cl^-^): Mean ± SEM: 19.4 ± 1.7 mV, Median: 21.3 mV, n = 15; Astrocyte: Mean ± SEM: 19.5 ± 3.1 mV, Median: 20.3 mV, n = 5; Ion-sensitive electrode: Mean ± SEM: 18.5 ± 2.4 mV, Median: 20.0 mV, n = 9. **(D)** During seizure-like activity, intracellular depolarizations and field potential fluctuations occur in a timely synchronous manner. **(E)** Representative recording of an astrocyte (left), pericyte (left center), endothelial cell (right center) and a human pericyte (right) exhibiting recurrent depolarizations either in the presence of 4AP or in Mg^2+^ free aCSF with elevated potassium and 10 μM Bicuculline for OHSC and human pericyte, respectively. Upper panels: morphological and electrophysiological properties of the recorded cell, lower panels: Vm changes with excerpt s showing the onset of a seizure. Red color in the first two pictures depicts MitoSox staining. Cells were either patched with Alexa 488 (green) or Alexa 568 (red) fluorescence dye.

### 5. Mechanisms underlying seizure-associated electrical signals in the capillary syncytium

While potassium determined the syncytial membrane potential, the observed depolarizations occurred from a negative resting potential, typical for pericytes in OHSCs [6]. In the presence of shear-stress and intramural pressure the membrane potential is more positive, and capillaries have a myogenic tone [45]. We previously used U46619 to reinstate capillary tone in OHSCs, in order to record seizure-dependent vasodilation [6]. We now investigated whether vasodilation in the presence of U46619 is accompanied by hyperpolarization of the capillary syncytium.

After switching to perfusion with 4AP the onset latency of seizures was 5-10 min, during which period neuronal network activity gradually increased [54] [51]. Pericytes hyperpolarized in the pre-seizure phase returning to values observed prior to U46619 application. As a result, the subsequent potassium-dependent depolarization during seizures began from a membrane potential of approximately −80 mV and never surpassed the level of depolarization initially induced by U46619 (Figure 5AB). In dual recordings of pericyte-pericyte or pericyte-endothelial cell pairs, both the pre-seizure hyperpolarization and the seizure associated depolarizations remained highly synchronous (Figure 5B). In summary, epileptic activity resulted in a sequence of hyper- and depolarization of the syncytium.

**FIGURE 5.**
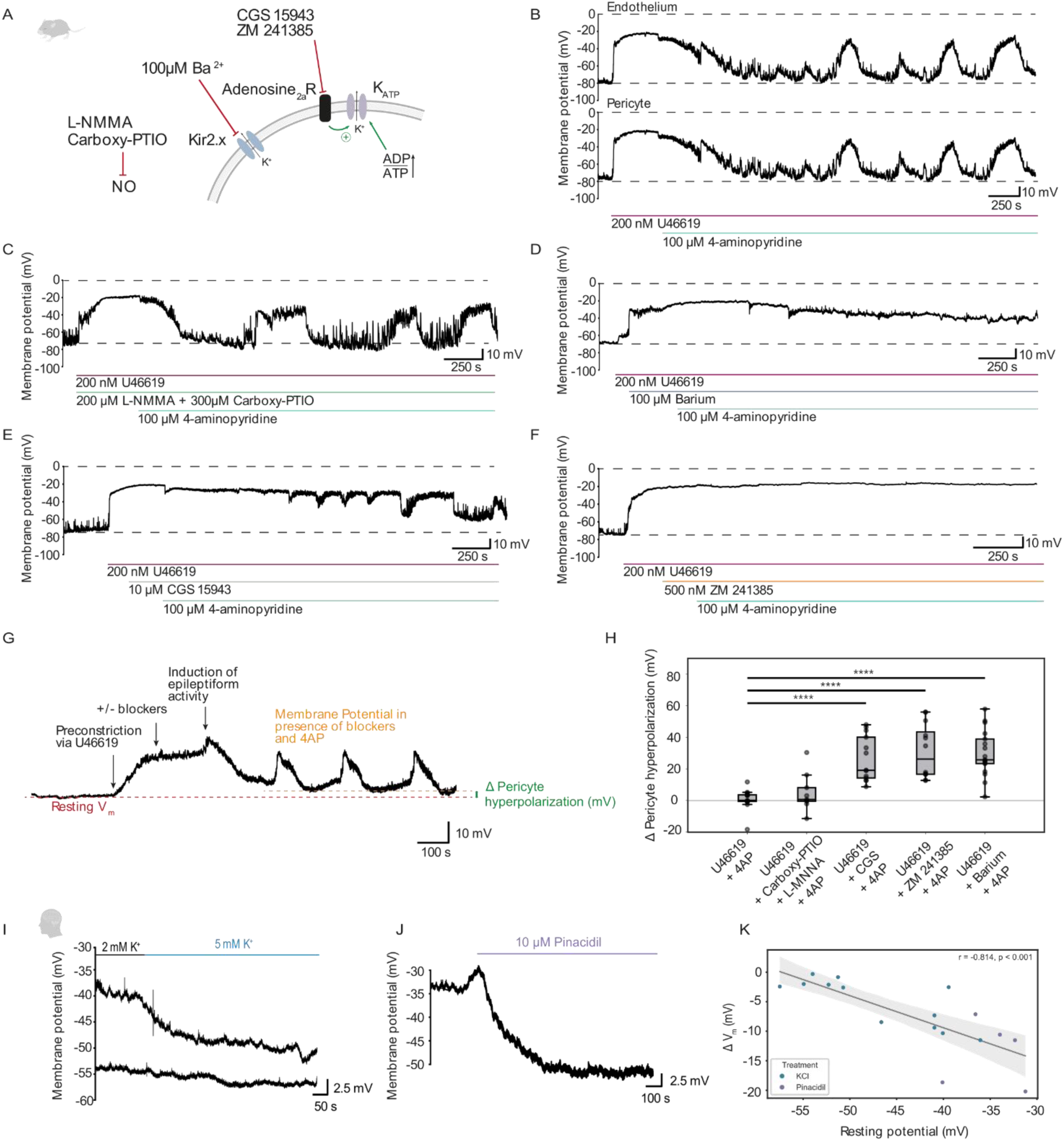
Seizure onset-induced hyperpolarization of pericytes via Kir 2.x channels and Adenosine 2a receptor (A2a) activation. **(A)** Schematic of the mode of action for the drugs used to interfere with pre-seizure hyperpolarization. **(B)** Representative dual-patch recording of a coupled pericyte-endothelial cell pair showing membrane potential changes during seizure-like activity in rat OHSC. The initial depolarization is caused by the thromboxane analogue U46619, which additionally constricts pericytes. At the onset of 4AP-induced seizure-like activity, the pericyte repolarizes to values observed prior to U46619, followed by recurrent seizure-associated depolarizations (dual patch: n=4, recordings from single pericytes n=15). **(C)** Hyperpolarization of pericytes could not be inhibited by application of a nitric oxide synthase inhibitor L-NMMA and the NO scavenger carboxy-PTIO (n=9). **(D, E, F)** Blocking Kir2.x channels using 100 μM barium (n=17), application of the non-selective adenosine receptor agonist CGS 15943 (n=13) or the selective A2a receptor antagonist ZM 241385 (n=12) partially or completely inhibited pericyte hyperpolarization. **(G)** Hyperpolarization of pericytes was determined as the difference between resting Vm prior to U46619 and the maximum of the pre-seizure hyperpolarization. Delta values close to zero represent a complete repolarization upon epileptiform activity. **(H)** Quantification of the repolarization in the presence of different blockers. Compared to U46619 + 4AP (U46619 + 4AP, p^ = 0.19 [0.15, 0.25]), significant differences were found for U46619 + 4AP and U46619 + CGS + 4AP (p^ = 0.64 [0.56, 0.72], p < 0.001), U46619 + 4AP and U46619 + ZM 241385 + 4AP (p^ = 0.70 [0.61, 0.77], p < 0.001), U46619 + 4AP and U46619 + Barium + 4AP (p^ = 0.69 [0.61, 076], p < 0.001). **(I)** In human cortical brain slices, pericytes hyperpolarize after an increase in potassium from 2 mM to 5 mM. **(J)** Hyperpolarization of pericytes after 10 μM pinacidil indicates the presence of KATP channels in human tissue. **(K)** The higher the resting potential of the pericyte before application of pinacidil (purple) and 3 mM potassium on top of 2mM aCSF K ^+^ concentration (blue), the stronger the hyperpolarization (Spearman correlation: ρ = -0.814, p < 0.001).

To identify the mechanism behind the pre-seizure hyperpolarization we pharmacologically targeted potential contributors: NO, potassium channels, and adenosine receptors. Neuronal NO formation is increased in the pre-seizure phase and contributes to seizure initiation in OHSCs [51]. As NO is a potent vasodilator, and NO-donors can reverse U46619-induced vasoconstriction [6] we next blocked NO-signaling by a combination of a NO-synthase inhibitor and NO-scavenger. Inhibiting NO-signaling had no effect on the pre-seizure hyperpolarization (Figure 5CI).

Endothelial cells and pericytes express inwardly rectifying potassium channels of Kir2.1, Kir2.2 and possibly Kir4.1 type. Perivascular potassium accumulation shifts the rectification of the channels to more positive values and increases their conductance, resulting in hyperpolarization of pericytes [27] [41]. Blocking Kir channels by 100 µM Barium chloride in the presence of U46619 and 4AP significantly delayed and decreased the amplitude of the pre-seizure hyperpolarization, indicating the contribution of these channels (Figure 5 DI).

Another vasoactive transmitter in control of ictogenesis is adenosine [55]; [56]. First, we used the potent adenosine receptor antagonist, CGS 15943 acting on A1, A2a, A2b and A3 receptors. CGS 15943 significantly decreased and delayed the pre-seizure hyperpolarization in pericytes (Figure 5EI). As A2a receptors are expressed on pericytes [27], we further refined the experiment using the selective A2a antagonist ZM 241385. Similarly to CGS 15943, ZM 241385 significantly decreased the pre-seizure hyperpolarization (Figure 5FI). A2a receptors may mediate hyperpolarization via Kir6.1, the most abundantly expressed potassium channel subunit in pericytes [29].

Next, we examined whether the mechanism of pre-seizure hyperpolarization in OHSCs also applies to human tissue. Upon perfusion with bicuculline-containing, low Mg²⁺ / elevated K⁺ aCSF, the human capillary syncytium hyperpolarized prior to the onset of seizure-like activity. This hyperpolarization largely depended on potassium concentration of the aCSF. In separate experiments increasing extracellular K⁺ from 2 to 5 mM in the absence of bicuculline and under normal Mg²⁺ conditions induced a similar hyperpolarization, supporting the involvement of Kir channels (Figure 5J).

Analogous to pre-seizure hyperpolarization in OHSCs, the K_ATP_ channel activator pinacidil lowered the syncytial membrane potential to -48.45 ± 2.95 mV (Figure 5K). The amplitude of hyperpolarization negatively correlated with the baseline membrane potential of the recorded cell (Figure 5L), suggesting that these channels were already active in a subset of cells. Thus, the mechanisms underlying pre-seizure hyperpolarization are conserved between rat and human capillaries.

## DISCUSSION

In this study, we provide evidence that mid-capillary mesh and thin-strand pericytes exhibit contractility and actively regulate capillary diameter. In the absence of blood flow and associated shear stress, local capillary tone was exclusively controlled by metabotropic receptor activation. Application of a thromboxane analogue, norepinephrine or UDP-glucose led to vasoconstriction and depolarization, regardless of the morphological class of the recorded pericyte. While depolarization and constriction happen simultaneously, the former did not contribute to vasoconstriction, as calcium influx via VGCCs was virtually absent in mid-capillary pericytes. Capillary diameter and pericyte length were unaffected by depolarizing voltage steps. Thus, the observed electrical signals may serve to communicate metabolic demands to ensheathing pericytes of the transition-zone and smooth muscle cells of upstream arterioles rather than to control the local capillary diameter. Indeed, in both slice cultures and human brain slices, the electrically coupled endothelial cells and pericytes acted as an “antenna,” recording and propagating neuronal activity-dependent changes in membrane potential. The electrical synapses between pericyte-pericyte and pericyte-endothelial cell pairs exhibited asymmetric coupling, a feature essential for the proposed upstream directionality of transmission. Prior to seizure onset, a potassium-dependent increase in Kir2.x channel conductance hyperpolarized the syncytium, even though extracellular potassium rises in the bulk parenchyma barely exceeded the detection limit of the potassium sensitive electrode. In addition to the Kir2.x channels, activation of A_2A_ receptors contributed to the pre-seizure hyperpolarization, indicating a dual mechanism for electro metabolic signaling. Despite its vasodilatory effect, seizure-associated nitric oxide formation had no significant influence on the capillary membrane potential. Subsequently, recurrent seizures repeatedly depolarized the syncytium reflecting changes in the Nernst potential for potassium.

### Heterogeneity of capillary pericytes

The term “pericyte,” as defined by Zimmerman (1923), refers to a heterogeneous group of mural cells associated with capillaries, initially classified based on morphological features. While recent classifications focus on the expression of specific surface markers, receptors, and ion channels, such as NG2, PDGFRβ, vimentin, and Kir6.1 among others, none of these markers are exclusively expressed by pericytes, and their expression varies along the arterio-venous axis [34] [27]. In this study, we utilized MitoSox, a mitochondrially targeted free radical marker [6] and NeuroTrace, used for Nissl staining [48], to label pericytes. Both probes labeled the same population of mural cells, and their specificity was confirmed in slices from transgenic animals [6]. Although slice cultures are prepared at a time (P6-8) when capillary development had not yet reached the adult pattern [57], superficial capillaries maintained an organotypic distribution. Due to the slicing, the branching order of individual capillary sections is often difficult to discern, however, neither of the probes labeled mural cells with morphology resembling SMCs. While we cannot completely exclude that the morphology of some NeuroTrace/MitoSox positive cells has been changed due to trans-differentiation under culture conditions, they still retained their contractility, electrical coupling and position at the capillary wall. The terminology of Grant et al. [36], classifies pericytes into ensheathing, mesh, and thin-strand subtypes, each likely contributing differently to blood flow regulation. In our study, only a very small fraction of fluorescent cells displayed the morphology of ensheathing pericytes, typical for post-arteriolar transition zone pericytes. Despite their morphological diversity, all subtypes, including thin-strand pericytes, exhibited constriction and depolarization in response to GPCR activation, likely with contribution of CaCCs.

Remarkably, the electrophysiological properties of the pericytes were not correlated with any of the morphological parameters. This is in line with previous findings in acute brain slices, where electrophysiological properties were homogeneous across morphologically distinct pericyte classes [41]. Finally, despite clear differences in cell size and resting membrane potential, human pericytes in acute slices displayed similar morphological and electrophysiological classes as those observed in slice cultures.

### Endothelial cell pericyte syncytium

Gap junctional coupling along the endothelial tube as well as between endothelium and adjacent pericytes is known in different organ vascular beds [58] [59] [60] [61] [62]. Besides the transport of small molecules, the relevance of gap-junctional coupling between endothelial cells lies in the transmission of locally generated electrical signals to distant feeding arterioles, termed the electro-metabolic signaling [14]. Coupling coefficient was remarkably high in the endothelial tube of isolated retinal vessels, and it dropped at the transition to precapillary arterioles, rendering these as an integrator for electrical signals [60]. Thus, overall increase in the supply by the arteriole necessitates a summation of dilatory electrical signals from several capillaries, whereas local capillary dilation might redistribute blood flow between adjacent capillaries [24].

Both in slice cultures and in human tissue all classes of recorded pericytes were coupled to the capillary syncytium, albeit the coupling index varied considerably. Dye diffusion from pericyte into adjacent endothelial cells always preceded staining of other pericytes even if their processes were in contact. At present we cannot distinguish whether this is due to a low efficiency of a direct coupling between pericyte pairs or a consequence of back-staining of pericytes via the endothelium.

In our paired recordings we found syncytial isopotentiality and a nearly simultaneous response to GPCR activation and extracellular potassium accumulation regardless of the pericyte morphology. While the electrical coupling between endothelial cells and pericytes was asymmetric and directed towards the latter in slice cultures, in human brain slices the direction was the opposite in half of the recorded pairs. Such asymmetric transmission might be brought about by differences in Rm of the individual cells or by the different composition of connexin subunits forming the gap junction [63]. As potential candidates for heteromeric gap junction, the expression of Cx37, Cx40, Cx43 and Cx30 have been described in pericytes and endothelial cells at different stages of the vessel maturation [64]. Electrical coupling coefficient was lower and asymmetric between pericyte-pericyte pairs, both in rat and human capillaries. Coupling asymmetry might underlie upstream directionality of hyperpolarizing signals towards transition zone capillaries [65] [15] [16] [66]. However, in a recent study potassium or glibenclamide induced hyperpolarization seemed to spread equally to up- and downstream vascular segments [67]. Thus, there might be additional pathways acting in concert which allow for dilation in upstream direction based on e.g. purinergic [68] [69] or nitrergic signaling [61].

We found significant differences between the syncytial membrane potential in acute slices and slice cultures. Pericytic membrane potential in pressurized retinal wholemount preparation was more positive (−40 mV) than in OHSCs, implicating the activation of mechanosensitive Piezo1 or transient receptor potential canonical (TRPC) channels [45] [70]. Although pressure and shear stress are completely absent both in slice cultures and in acute brain slices, Vm was still fairly positive in the latter. One possible explanation is that cutting capillaries during slicing inevitably exposes them to damage-associated molecular patterns (DMPPs), such as endothelin-1, thromboxane, ATP and UDP-glucose. Vasoconstriction and depolarization of the pericytes is a ubiquitous response upon injury. Indeed, pericytes were depolarized in both rat and human acute brain slices, suggesting that interspecies differences would not contribute to the finding. During the course of the first two days in culture the capillary syncytium hyperpolarizes in the absence of mechanical forces and stabilizes at ∼ -80 mV, for up to two weeks in vitro. While we cannot exclude that changes in ion channel expression might contribute to the hyperpolarized Vm in culture, it is worth to note that mechanical activation still induces depolarization and constriction of pericytes, indicating the presence of mechanosensitive Ca^2+^ influx even after 14 DIV ([6], supplemental figure).

### Electro-metabolic coupling and capillary vasomotility

The hypothesis of vascular electro-metabolic signaling [14] postulates that metabolic alterations in the tissue lead to changes in vascular syncytial membrane potential, which - in turn- are responsible for setting the vasotonus. In line with this hypothesis, we found that activation of TX2A receptors simultaneously depolarizes mid-capillary pericytes and triggers vasoconstriction. In SMCs, Gq coupled TX2A receptors trigger inositol 1,4,5-trisphosphate (IP₃)- and ryanodine receptor (RyR)-mediated Ca²⁺ release from intracellular stores. The subsequent opening of CaCCs depolarizes SMCs ultimately leading to additional Ca²⁺ influx through VGCCs and further amplifying vasoconstriction [71]. Indeed, large pathological depolarizations as a consequence of hypoxia or spreading depolarization can result in Ca^2+^ overload in SMCs and in ensheathing pericytes. This mechanism is expected to contribute to the no-reflow phenomenon in the brain, heart and kidney [72] [73] [74] [75] [46]. While electro-metabolic coupling is primarily mediated by the activation of VGCCs in SMCs and in pericytes of the transition zone, in mid-capillary pericytes store-operated calcium entry (SOCE) and TRPC channels are responsible for Ca^2+^ influx, while voltage dependent sources seem less relevant [44] [45] [76] [77]. Indeed, Gonzales et al. [24] demonstrated that mid-capillary pericytes contract in response to U46619, but not in response to depolarizing concentrations of extracellular potassium (60 mM).

Here, we found further evidence that constriction of mid-capillary pericytes is independent of the membrane depolarization. Different depolarizing protocols via the patch-pipette failed to elicit voltage gated Ca^2+^ currents, to alter intracellular Ca^2+^ concentration or induce pericyte length changes. Even pharmacological activation of VGCCs by BAY-K-8644 resulted in only a slow and delayed intracellular Ca^2+^ increase in a subset of cells, suggesting that any VGCCs present, are usually inactive under culture conditions [30] [78].

In conclusion, local GPCR mediated capillary constriction appears independent of the associated depolarization within the capillary syncytium. Thus, the latter might be more relevant to upstream targets than the capillary itself. On the contrary, mechanical forces unequivocally depolarize and constrict mid-capillary pericytes indicating the presence of mechanosensitive Ca^2+^ influx pathways ([6], supplemental figure).

### Epileptiform activity induced changes in the syncytial potential

In a previous study we have shown that during recurrent epileptiform activity each individual seizure was associated with a vasodilation [6]. Regarding electrical signaling, here we found that capillary syncytium hyperpolarizes even before seizure onset despite the continued presence of U46619, while seizures resulted in positive shifts in the membrane potential.

One plausible explanation for the pre-seizure hyperpolarization is offered by neuronal NO release, which controls the basal tone of transition zone pericytes [68]. Previously we found that neuronal nitric oxide synthase activity increases before the onset of the first seizure [51], and NO-donors dilate pre-constricted capillaries in cultures [6]. However, despite the vasodilatory effect, the blockade of NO signaling did not prevent the pre-seizure hyperpolarization, indicating that the observed dilation was not dependent on membrane potential.

Another potential candidate is the abundantly expressed KATP channel, whose pharmacological activation induces capillary dilation in different models [68] [79] [67] [80]. In our hands, inhibition of A2a receptors significantly decreased the pre-seizure hyperpolarization, substantiating a role of K_ATP_ channels. Gαs coupled A2a receptors stimulate the synthesis of cAMP, which in turn leads to PKA dependent phosphorylation and opening of the K_ATP_ channel complex. Consequently, A2a agonists were able to prevent capillary pericyte constriction and cognitive impairment in a model of chronic cerebral hypoperfusion [81]. On the contrary, activation of Gαi coupled P2Y14 receptors by UDP-glucose resulted in a depolarization, further underscoring the relevance of Kir6.1 channels in syncytial potential regulation. While significant extracellular accumulation of adenosine typically occurs only during seizures [81] [56], our results suggest that capillaries are exposed to adenosine early in the seizure initiation process.

Nevertheless, in most slices, A2a receptor blockade did not fully prevent the pre-seizure hyperpolarization, indicating the involvement of additional mechanisms. Potassium dependent activation of endothelial Kir2.1 channels generate propagating hyperpolarizing signals to dilate parenchymal arterioles [23] [29]. Consistent with this hypothesis, application of Barium chloride partially prevented the pre-seizure hyperpolarization, suggesting endothelial and pericytic Kir2.x channel involvement. The hyperpolarization mediated by the A2a receptor might relieve the Mg^2+^ / polyamine block of the Kir2.x channels thereby fostering the spread of hyperpolarization and preventing activation of VGCCs in upstream transition zone pericytes.

This is intriguing in light of previous studies showing dramatic, VGCC-dependent prolonged arteriolar vasoconstriction following seizures [4]. Our results suggest that postictal vasoconstriction at the level of higher order capillaries is independent of voltage gated Ca^2+^ influx despite the repeated depolarization. Alternatively, metabolic disturbances might underly the lasting postictal constriction as signs of mitochondrial dysfunction were present in constricting pericytes [6]. In the current experimental setting—where ATP was abundantly supplied via the pipette solution—pericytes exhibited increased resistance to recurrent seizures lasting over 40 minutes. The precise mechanisms by which seizures induce metabolic disturbances in pericytes remain to be elucidated.

## MATERIALS AND METHODS

### Study approval

Animal experimental protocols were approved by the Animal Ethics Committee of the Charité – Universitätsmedizin Berlin. Animal care and handling was in accordance with the WMA Declaration of Helsinki and institutional guidelines (https://experimentelle-medizin.charite.de/en/) as reviewed by the Berlin State Office for Health and Social Affairs (T-CH 0003/20). The conduct of experiments with human tissue from neurosurgical patients was approved by three independent ethics committees (Charité - Universitätsmedizin Berlin: vote no. EA2/111/14, EA2/064/22 in reference to EA4/206/20, EA2/086/20; Ärztekammer Westfalen-Lippe und der Westfälischen Wilhelms-Universität: vote no. 2020-517-f-S; Ärztekammer Hamburg: vote no. 2023-200674-BO-bet in reference to EA2/111/14). Written informed consent was obtained from all patients for the scientific use of their resected tissue. This study was reported in accordance with the MDAR guideline [82].

### Materials availability

For the purpose of this study no new reagents were developed.

### Organotypic slice culture preparation and staining

Organotypic hippocampal slice cultures (OHSC) were prepared and cultured according to Stoppini’s culturing method [83] [47]. The hippocampi of P6 to P8 wistar rat pups of either sex were cut into 400 μm thin slices under ice-cold, carbogen-gassed (95 % O_2_, 5 % CO_2_, pH 7.3) minimal essential medium (MEM), placed on a culture plate insert (MilliCell-CM, Millipore, Eschborn, Germany) and kept in an incubator for 5-12 days in vitro (DIV) before using in experiments. Culture medium (50 % MEM, 25 % Hank’s Balanced Salt Solution, 25 % Horse Serum, pH 7.4; Gibco, Eggenstein, Germany) was substituted three times a week. OHSCs were stained in the incubator at 37 °C with either the mitochondrially targeted ethidium derivative MitoSOX (Invitrogen, 5 μM, dissolved in DMSO, 0.1 % in final solution) or with the NeuroTrace 500/525 Nissl stain (Invitrogen, dissolved in DMSO, 0.1 % final solution) >20 min prior to the experiments.

### Human brain tissue and preparation of human acute slices

The human temporal and frontal association cortical tissue was obtained from neurosurgical patients undergoing surgery to treat drug-resistant epilepsy. The age of the patients ranged from 3 to 58 years (mean: 30.26 ± 2.73 years, Table 1). Resection and slicing followed the procedures described in [84]. In brief: resected tissue was immediately placed in sterile, ice-cold, carbogen-saturated, sucrose-based artificial cerebrospinal fluid (saCSF) comprised (in mM): 87 NaCl, 1.25 NaH_2_PO_4_, 2.5 KCl, 0.5 CaCl_2_, 3 MgCl_2_, 10 Glucose, 25 NaHCO_3_, 75 Sucrose, pH 7.4, 310 mOsm/l. Following transport to the laboratory, pia mater was removed, tissue block trimmed and cut into 300 μm-thick slices perpendicular with the cortical surface. After a 30-minute recovery period at 34 - 36 °C, the slices were cooled down and kept at room temperature (22 – 24 °C) in sterile saCSF until used for experiments 2 - 48 hrs. Before the patch clamp recordings, each slice was digested for 20 minutes at 36°C with protease XIV (0.4 mg/ml; Sigma-Aldrich) and collagenase type I (0.5 mg/ml; Rockland) in human aCSF containing (in mM): 125 NaCl, 1.25 NaH_2_PO_4_, 2.5 KCl, 2 CaCl_2_, 1 MgCl_2_, 10 Glucose, 25 NaHCO_3_, 1 ascorbic acid, pH 7.4, 300 mOsm/l).

**Table 1.** xxxxx

|  |  |
| --- | --- |
| Patient age | 30.26 ± 2.73 years |
| Age of onset | 16.72 ± 2.21 years |
| Duration | 13.84 ± 2.22 years |
| Sex | female: 15, male: 20 |
| Hemisphere | Left: 20 right: 15 |
| Structure/Lobe | temporal: 30, frontal: 3, occipital: 1, hemispherectomy: 1 |
| Pathology | Hippocampal sclerosis: 16<br>Gliosis: 3<br>FCD/MOGHE: 5<br>Polymicrogyria: 2<br>Meningo/encephalocele: 3<br>Cavernoma: 1<br>Ganglioma: 3<br>Rasmussen/Hippocampal encephalitis: 2 |

### Electrophysiological recordings

Human cortex slices and rat slice cultures were transferred to a recording chamber perfused either with carbogen bubbled human aCSF (as above) or aCSF for slice cultures (containing 129 NaCl, 10 glucose, 1.25 NaH_2_PO_4_, 1.6 CaCl_2_, 3 KCl, 1.8 MgSO_4_ and 26 NaHCO_3_ 310–315 mOsmol/kg and a pH of 7.4) at a perfusion speed of 5 ml/min (31 to 32°C). To improve oxygen availability and foster network activity a double-perfusion recording chamber [85] was used for thick human slices.

As NeuroTrace staining functions less well in human brain slices, capillary pericytes were identified here by their typical “bump-on-a-log” appearance at small vessels with a diameter of < 10 µm. Their identity was further proved by reconstruction following staining via the pipette and also by the presence of electrical coupling to other components of the capillary syncytium, which is not present in other perivascular cell types, such as OPCs, microglia or myofibroblasts. In contrast, accumulation of NeuroNissl or oxidized MitoSOX was reliable - and overlapping - marker of pericytes in OHSCs. Patch pipettes with a tip resistance ∼ 8-12 MΩ were made from borosilicate glass capillaries (Science Products, Hofheim am Taunus, Germany). We used three different intracellular solutions (i) a low chloride potassium-gluconate solution, consisting of (in mM): 130 K-gluconate, 4 KCl, 2 MgCl_2_, 4 Na_2_-ATP, 5 Na_2_-phosphocreatine, 0.3 Na-GTP, 10 HEPES or (ii) a high chloride potassium-gluconate solution, consisting of (in mM): 105 K-gluconate, 30 KCl, 2 MgCl_2_, 10 HEPES, 4 Na_2_-ATP, 5 Na_2_-phosphocreatine, 0.3 Na-GTP (pH set with 420 µl KOH to 7.2, Osmolarity: 282 mOsmol/kg) (iii) and a CsCl-based intracellular solution (in mM): 135 CsCl, 3 NaCl, 0.1 CaCl_2_, 1 EGTA, 10 HEPES, 2 Mg-ATP (pH set to 7.25 with KOH/CsCl, osmolarity ∼275-290 mOsmol/kg). For experiments with dye coupling and calcium imaging either Alexa Fluor 488 (100 µM), Alexa Fluor 568 (100 µM), 0.3125 Oregon-Green Bapta-1 hexapotassium (Kd = 170 nM) or 0.5 rhod-2 tripotassium (Kd = 570 nM) was added to the intracellular solution. Whole-cell recordings were made using a Multiclamp 700B amplifier and pClamp 10 software (Axon CNS, Molecular Devices). Data were low-pass filtered with an 8-pole Bessel filter at 3 kHz and digitized at 10 kHz using Digidata 1440A (Axon CNS, Molecular Devices). The presented values were not corrected for liquid junction potential.

### Local field potential and ion-sensitive electrode recordings

Measurements of local field potentials (fp) were made in the stratum radiatum of CA3 in OHSCs with a glass electrode filled with aCSF. Changes in ([K^+^]o) were determined using double-barreled ion-sensitive microelectrodes prepared and tested as previously described [51]. The potassium concentration is calculated from the potential of the ion-sensitive side of the electrode by a modified Nernst equation, while neuronal network activity is recorded as a field potential at the reference side. The reference cylinder was filled with 154 mM NaCl solution, the ion-sensitive cylinder with 100 mM KCl and the tip with the ionophore cocktail potassium ionophore I 60031 (Fluka, Buchs, Switzerland). The signals of the home -built ion-sensitive amplifier were filtered at 300 Hz and recorded at 1 kHz by using Power1401 and Spike2 software (Cambridge Electronic Design Limited, Cambridge, UK).

### Confocal microscopy and Ca^2+^ Imaging

3D reconstructions and calcium imaging of the recorded cells and capillaries were acquired using a spinning-disc confocal microscope with 60x objective (NA = 1.1) and the Andor IQ software (Andor Revolution, BFIOptilas GmbH, Gröbenzell, Germany). To monitor changes in intracellular Ca^2+^ concentration fluorescent images were acquired either as time series in a single plane (7.7 to 8.5 frames per second) or as repeated z-scans spanning the entire capillary every 10 - 20 s for >10 min in case of dye coupling. Simultaneous recordings in multiple capillaries were obtained with a NIKON A1R MP multiphoton microscope (25x N.A. 1.1 objective, Nikon, Amsterdam, The Netherlands) at the AMBIO Life Cell Imaging Core Facility (AMBIO.charite.de).

### Drugs

To constrict capillaries UDP-glucose (100 µM, Merck), norepinephrine (10 µM, Sigma A7256) and the thromboxane agonist, U46619 (Stock solution: 2 mM, Tocris bioscience, diluted in DMSO) were applied via the perfusion. Epileptiform activity was induced by using the voltage-gated potassium channel inhibitor 4-aminopyridine (4AP; Stock solution: 100 mM, Tocris bioscience, diluted in H_2_O) in OHSCs and bicuculline methiodide (Stock solution: 10 mM, Research Biochemicals International, diluted in DMSO) in combination with Mg^2+^ free aCSF containing 87 NaCl, 1.25 NaH2PO_4_, 5 KCl, 0.5 CaCl_2_, 10 Glucose, 25 NaHCO_3_, 75 Sucrose, 1 ascorbic acid in human brain slices. Stereotypic ictal discharges in local field potential resembling seizure activity, i.e. seizure-like events, are referred to as “seizures” throughout the text.

To interfere with the pre-seizure hyperpolarization CGS 15943 (Stock solution: 20 mM, Cayman Chemical Company, diluted in DMSO), ZM 241385 (Stock solution: 10 mM, Tocris Bioscience, diluted in DMSO), Barium chloride (Stock solution: 100 mM Barium-aCSF, Sigma), Carboxy-PTIO (Stock solution: 3 M, Alexis Biochemicals, diluted in aCSF) and L-NMMA (Stock solution: 200 mM, Alexis Biochemicals, diluted in aCSF) were applied several minutes before 4-AP wash in. To activate VGCCs we used BAY-K 8644 (Stock solution: 1 mM, final 100nM, Cayman Chemical Company, diluted in DMSO).

### Image and Morphological Analysis

Image analysis for time series recording was performed using either the NeuroOptical Signal Analysis (NOSA) software [86] or ImageJ [87]. Motion artifacts were reduced either by symmetric diffeomorphic registration or by rigid body transformation (NOSA) and the baseline fluorescence was corrected for photobleaching and signal drift. Fluorescence signals of selected regions of interest (ROI) were smoothed by applying a Savitzky-Golay filter. Following background subtraction fluorescent transients in Ca^2+^-imaging were presented as ΔF/F_0_ (where F_0_ originates from the ROI prior to voltage steps or drug application. At the end of patch clamp recordings, higher resolution z-stack images of the pericyte were obtained for morphological and dye coupling analysis. Coupling index was defined as the ratio of fluorescence of the processes of the recorded cell and subsequent cells of the syncytium after reaching steady state in dye diffusion. Z-stack images were pre-processed using Gaussian blur (σ = 1) and rolling ball background subtraction (radius = 25 pixels). For cellular segmentation Li’s automatic thresholding method was used and local thresholds adjusted manually to ensure that the whole pericyte is presented including thin processes. Three-dimensional morphological parameters were calculated using MorphoLibJ [88] and Analyze Skeleton [89] plugins.

### Analysis of the electrophysiological data

Analysis and visualization of the electrophysiological data were performed using custom-written Python scripts (Python 3.12, pyABF package: [90, 91]). Series (Rs), input resistance (Ri) and current to voltage relationship (from -100 to +50 mV in 10 mV steps) was determined immediately after establishing the whole-cell configuration. Rs and Riwere calculated using a 10 mV voltage step by automatically extracting baseline, maximum and steady-state values from each sweep (Rs OHSC: Mean ± SEM: 25.55 ± 0.677 MΩ, n = 323, Rs human: Mean ± SEM: 25.05 ± 0.87 MΩ, n = 93). Ri is given after being corrected for Rs but includes all electrically coupled components of the syncytium. Rm was not assessed in cells where gap junctions were pharmacologically blocked to isolate individual cells. The resting membrane potential (Vm) was determined using the median zero-current potential before switching from voltage to current clamp for recording GPCR and activity dependent changes in Vm. In dual patch recordings several trials of de- and hyperpolarizing current pulses (−100, -50, +50 pA) were given alternating to each cell, and the averaged voltage responses from the stimulated and receiver cell were divided to determine the coupling coefficient in each direction.

### Statistical analysis

The statistical analysis was performed using R (Posit PBC, R Version 4.4.2). Due to small sample sizes and skewed data distribution, nonparametric tests were used (nparcomp, [93]). Nonparametric multiple contrast tests were used to compare between-group differences across all experimental conditions. Dunnett nonparametric multiple contrast tests were used to compare the membrane potential of pericytes in different preparations as well as the pre-seizure pericyte repolarization in the presence or absence of blockers. A Tukey nonparametric multiple contrast test was used to test all pairwise differences in the depolarization amplitude upon U46619 application between the three experimental groups. Systematic differences between paired measurements were assessed using the nonparametric studentized permutation test for paired data. The permutation-based p-value (PERM) was reported throughout. P-values from multiple contrast tests were multiplicity-adjusted. For multiple contrast tests (mctp), results are reported as marginal relative effects (p^) with 95% confidence interval (CI). For paired comparisons (npar.t.test.paired), results are reported as pairwise relative effects (p^) with 95% CI. Where confidence intervals exceeded the theoretical bounds of [0, 1], they were capped accordingly. Spearman’s rank correlation was used to test the strength of associations between membrane resistance and coupling coefficient as well as initial resting membrane potential before and after application of pinacidil or 5 mM potassium. All tests were two-sided. P-values ≤ 0.05 were considered statistically significant. Data are reported as mean ± SEM or median and interquartile range (IQR). Statistical advice was provided by the Biometric Institute of the Charité.

### Cluster analysis

We performed two separate cluster analyses for morphological and electrophysiological parameters since the initial attempt to perform combined clustering of all parameters together failed to produce distinct and well-defined clusters. Therefore, we decided to cluster both attributes separately. Cluster analysis was carried out to identify pericyte subtypes based on morphological features within the population of patched capillary pericytes (n = 336). Considering known pericyte types, we initially selected four features with high discriminatory power: (1) ’Longest Shortest Path’ as a representation of the longitudinal axis of the cell, (2) ’Elli.R1/R3’ the ratio of the encircling ellipsoids as a parameter for directional elongation and spindle-shape forms, (3) sphericity to capture more stellate morphologies, and (4) the ’number of junctions’ correlating with number of processes building the “mesh”. In a subsequent step, the silhouette score was used to test whether the addition of further parameters such as the ’Number of end-point voxels’, ’Surface Area divided by Volume Ratio’, ’Average Branch Length’ or ’Raw voxel count’ could increase the discriminatory power of the groups. The four selected features resulted in a silhouette score of 0.29 for four clusters while adding further parameters did not significantly increase the silhouette score. The elbow method supported the decision of four groups. A separate cluster analysis was performed for the electrophysiological features. This included the current-voltage relationships (IV), membrane resistance (Ri), and resting membrane potential (Vm). The current-to- voltage curve was determined for the steady-state current at the end of each current step and each IV curve is presented after normalizing to its maximum current. To obtain characteristic variables describing of the shape of the IV curves, a 4th-degree polynomial fitting (I(V) = a₄V⁴ + a₃V³ + a₂V² + a₁V + a₀) was performed using least-squares regression (numpy.polyfit) on each normalized IV curve. As a metric of rectification, we selected the third coefficient (a₂), since a higher absolute value of a₂ indicates greater deviation from linearity. Cells that showed strong IV outliers were removed before the clustering process (culture: 13 of 336, human tissue: 4 of 97 outlier IV curves), based on the assumption that recording configuration did not meet quality criteria (bad seal, current leak). Outliers of the membrane resistance values determined using the 1st and 99th percentile as threshold values. Values outside this range were replaced with the respective boundary values via winsorization. Prior to clustering, features were standardized using scikit-learn’s standard scaler (scikit-learn version 1.4.1., [92]). Due to 6.23% missing data points, clustering was performed using KmeansWithNulls (version 0.1.1). The debugging and development of Python and R code was supported by a large language model (Claude Sonnet 3.5 and 4.6, Anthropic). All code was reviewed and validated by the authors, who take full responsibility for their accuracy and interpretation. The AI tool was not used for data interpretation, generation of scientific conclusions, or manuscript writing beyond code-related tasks.

## Code availability

All code used for data analysis and figure generation is publicly available on GitHub at https://github.com/mgrotelambers/pericyte-syncytial-coupling-seizure-signaling.

## Data availability

The data that support the findings of this study are available on request from the corresponding author. The data are not publicly available due to privacy or ethical restrictions.

## ACKNOWLEDGEMENTS

This study received support from the DFG grant CRC 1365, which funded RK. MGL received a doctoral scholarship from the Charité. The authors extend their heartfelt gratitude to Mrs. Andrea Wilke for her invaluable technical assistance, Prof. Alon Friedman for fruitful discussions and would like to thank Stephen Schüürhuis for his valuable statistical advice.

## AUTHOR CONTRIBUTIONS

RK, MGL, CM, and JRPG conceptualized and designed the project. MGL, HW, HP, and RK performed electrophysiological data acquisition and imaging. AL conducted potassium recordings. MK wrote the ImageJ macro code for the image analysis. MGL performed the image analysis. MGL carried out the analysis of electrophysiological data, statistics and data visualization. TK, RX, JO, TS, UWT, AK, MH and MS contributed to neurosurgical tissue acquisition and patient care. PF managed the REDCap patient database and coordinated patient data collection. PF, HA and JRP handled institutional collaborations and ethical approvals. MGL, RK, CM, and JRPG wrote the original draft of the manuscript. All authors reviewed and edited the manuscript.

## DECLARATION OF INTERESTS

The authors declare no conflict of interest.

## Notes

### Competing Interest Statement

The authors have declared no competing interest.

### Summary of Updates

We have corrected small typing errors, added new statistics (marginal relative effects (p&#770;) and pairwise relative effects (p&#770;) with 95% CI.), added a table with patient demographic/pathological data

