## Supplemental Figure 1 for "Syncytial coupling of mid-capillary pericytes underlies seizure-associated electro-metabolic signaling"

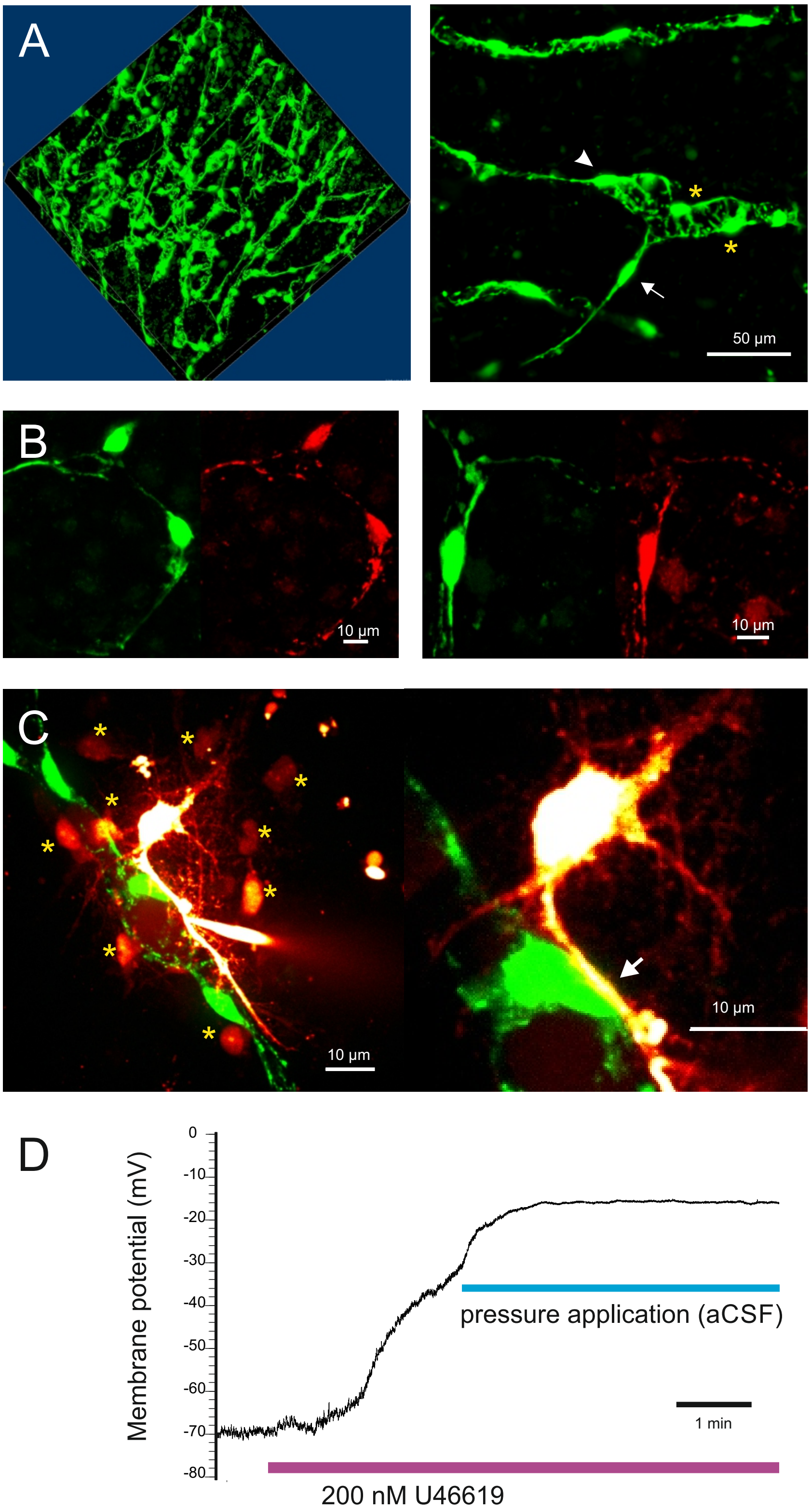


**(A)** Distribution of NeuroTrace stained capillaries in hippocampal slice cultures. Left: Slightly tilted 3D reconstruction of the vasculature at the CA3-CA1 border in the stratum radiatum. Due to slicing and the density of the capillary mesh, exact branching order is difficult to ascertain in cultures. Right: Maximum intensity projection of capillary bifurcation showing thin-strand- (arrow), mesh- (asterisk) and transitional pericytes (arrowhead). **(B)** Co-labeling of slice cultures with MitoSox (red) and NeuroTrace (green). While the unspecific ROS indicator MitoSox showed mitochondrial labeling throughout the parenchyma and sometimes in neuronal somata, pericytes were unequivocally co-labeled with both probes. **(C)** Single cell recordings with rhod-2 containing pipette solution obtained at the astrocyte endfeet revealed high coupling index with neighboring astrocytes (asterisks) while the underlying pericytes remained unstained despite the close contact (right, arrow). **(D)** Pressure induced pericytic depolarization. Following U46619-induced depolarization of the pericyte, pressurized aCSF flow from a nearby pipette targeting the recorded cell still evoked an additional depolarization, indicating the presence of mechanosensitive cation channels.
